# Humans could become the greatest driver of biosphere net gain in Earth history, but are currently the second fastest driver of biosphere loss

**DOI:** 10.64898/2026.04.10.715592

**Authors:** Thomas W. Wong Hearing, Mark Williams, Jan Zalasiewicz, Heiko Balzter, Davor Vidas, John Maltby, Julia Adeney Thomas, Sergei Petrovskii, Colin N. Waters, Martin J. Head, Libby Robin, Elizabeth Hadly, James Borrell, Colin Summerhayes, Alejandro Cearreta, Anthony Barnosky, Francine McCarthy, Pat Heslop-Harrison, Reinhold Leinfelder, Sverker Sörlin, Jens Zinke, Michael Wagreich, Moriaki Yasuhara

**Affiliations:** School of Geography, Geology and the Environment, University of Leicester, UK; Institute for Environmental Futures, University of Leicester, UK; National Centre for Earth Observation, Space Park Leicester, UK; Fridtjof Nansen Institute, Oslo, Norway; School of Healthcare, University of Leicester, UK; Department of History, University of Notre Dame, Indiana, USA; School of Computing and Mathematical Sciences, University of Leicester, UK; Department of Earth Sciences, Brock University, St Catharines, Ontario, Canada; Fenner School of the Environment and Society, The Australian National University, Australia; Department of Biology, Stanford University, Stanford, USA; Royal Botanical Gardens Kew, Richmond, London, UK; Scott Polar Research Centre, University of Cambridge, Cambridge, UK; Departamento de Geología, Universidad del País Vasco UPV/EHU, Bilbao, Spain; Department of Integrative Biology, University of California, Berkeley, USA; Department of Genetics and Genome Biology, University of Leicester, UK; Department of Geological Sciences, Freie Universität Berlin, Germany; Division of History of Science, Technology and Environment, KTH Royal Institute of Technology, Stockholm, Sweden; Department of Geology, University of Vienna, Austria; School of Energy and Environment, State Key Laboratory of Marine Environmental Health, City University of Hong Kong, Hong Kong SAR, China

## Abstract

We evaluate episodes of biosphere change throughout Earth history and compare them with contemporary and near-future anthropogenic changes, developing the concept of *biosphere disruptors* – agents that force global-scale macroevolutionary change. *Transient disruptors* are geologically short-lived agents (mean 8.0×10^5^ years), including massive volcanism and asteroid impacts. *Persistent disruptors*, including atmospheric and ocean oxygenation and land plant evolution, remain in the Earth System over long timescales (mean 1.3×10^8^ years). In the geological record, *transient disruptors* are associated with temporary but sometimes massive biosphere degradation, whereas *persistent disruptors* are associated with sustained biosphere enhancement. Most anthropogenic biosphere impacts resemble those of past *transient disruptors*, globally degrading wild biomass and biodiversity. Humanity is driving the second highest rate of biosphere degradation in Earth history after the Cretaceous-Paleogene asteroid impact. However, humanity is the first disrupting agent capable of reflecting on and potentially transforming its impact on planetary habitability. If this can be achieved, humanity could drive the greatest rate of increase in planetary habitability in Earth history on centennial to millennial timescales.

## 1. Introduction

Seven of the nine planetary boundaries that determine a safe operating space for human interactions with Earth’s life support systems are transgressed by human activity (figure 1) [1]. The two components of the biosphere integrity boundary have been transgressed over the last 150 years [2,3]. Functional integrity, representing the biosphere’s capacity to maintain ecological functions, was breached in the 19^th^ century when human-appropriated net primary production (HANPP) first exceeded 10%; by 2023 HANPP had increased to ∼30% [3]. Genetic integrity follows extinction rate, with the boundary value of fewer than 10 extinctions per million species years (E/MSY) far exceeded in the last 150 years [2]. The current value exceeds 100 E/MSY [3], with ∼10% of genetic diversity lost since the 19^th^ century [1,4]. Population loss is typically a precursor to species extinction, and monitored vertebrate populations decreased by an average of 73% of their 1970 value by 2020 (<u>The Living Planet Index)</u>, with similar declines in invertebrate groups including insects [5].

**Figure 1.**
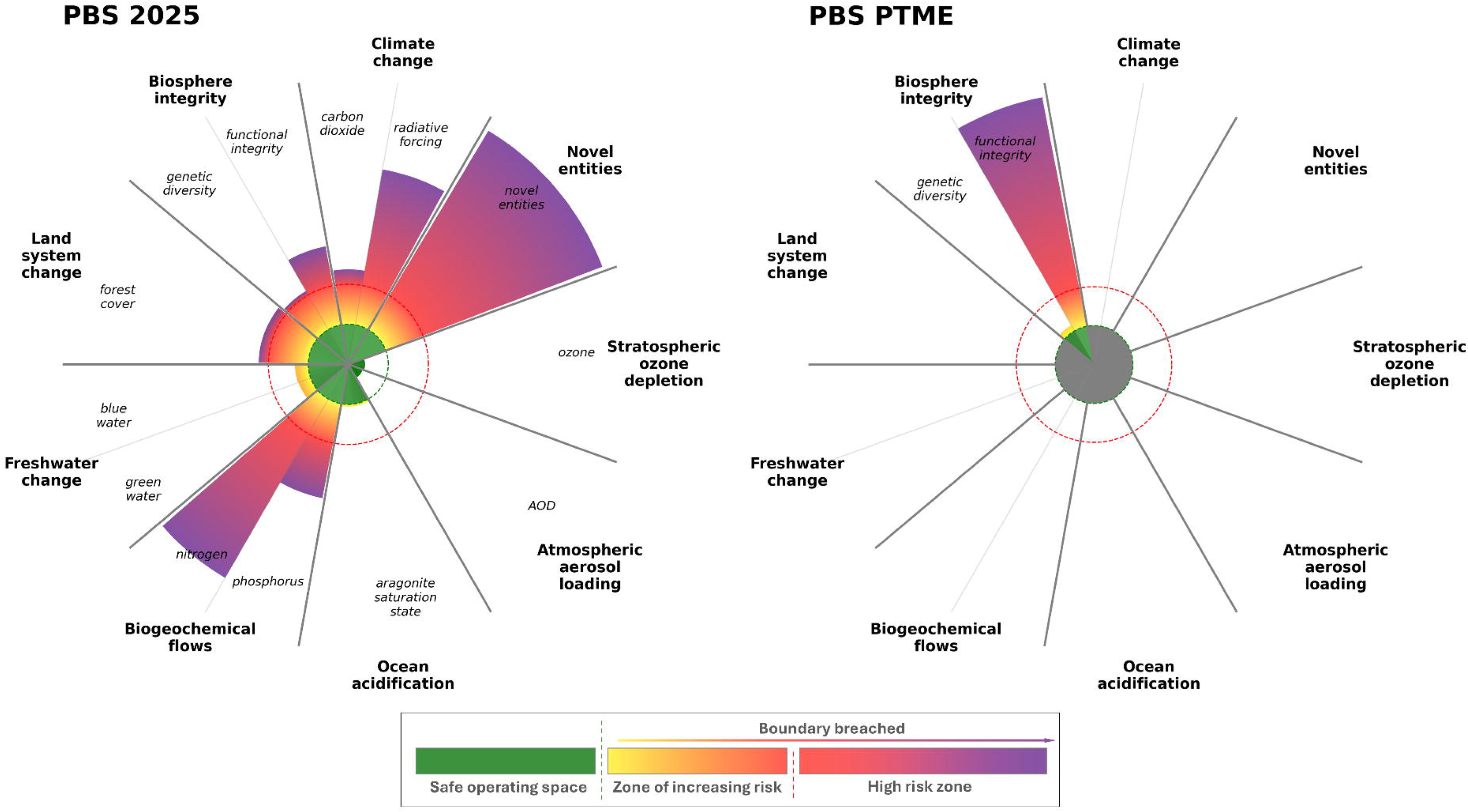
The planetary boundaries framework reveals that humans are pushing the Earth System beyond the safe limits of its recent past (PBS 2025; [1]). Compare the 2025 biosphere integrity boundary values with those of the Permian-Triassic Mass Extinction ∼252 million years ago (PBS PTME). The PTME species extinction value of ∼95% equates to ∼15 extinctions per million species years (see tables S3 and S5). Green dashed line: threshold for safe operating space; red dashed line: boundary of the zone of increasing risk and the high-risk zone. We have deliberately not attempted to quantify other elements of the PTME planetary boundaries, but we encourage further work applying the planetary boundaries framework to past intervals.

Human influence on planetary function increased rapidly during the mid-20^th^ century ‘Great Acceleration’ [6] and is continuing in the 21^st^ century [1,3]. Anthropogenic changes to Earth’s life support systems have been predominantly driven by humans in hyper-consuming societies, rather than humanity in total [7,8]. Today, most of Earth’s atmosphere, oceans, and terrestrial surface are pervasively influenced by humans [9,10]. Land area covered by primary vegetation has fallen from ∼70% in 1850 to <40% in 2000 [11], and the proportion of unsustainable fisheries increased from ∼10% in 1974 to ∼35% in 2021 (though the proportion of under-exploited stocks has increased sharply since 2019) [12].

Humans have approximately tripled total mammal biomass since 1850 through population growth and an ∼500% increase in domesticated livestock [13]. The cost of this has been the loss of ∼60% of terrestrial and ∼70% of marine wild mammal biomass [13]. Conservation can counteract some of these losses, and conservation efforts have halted and substantially reversed some regional population declines, including range expansions of some species [14]. Some land management practices also benefit both human societies and the local wild biosphere [15], including matrix farming (e.g. [16]), patch-burning (e.g. [17]), and sustainable fishing [18]. Technological developments also provide space for the evolution of novel ecologies (but see [19]), and some local biodiversity increases have been driven by deliberate or accidental anthropogenic species translocation and hybridization (e.g. [20]) and probably by the allopatric speciation of non-native species isolated from their native gene pool [21].

We suggest that lessons drawn from large biotic changes in Earth history can inform the actions necessary for humanity to augment planetary habitability. Here, we use planetary habitability in the sense of Earth’s capacity to support all of functional and taxonomic biodiversity, productivity, biomass, and resilience across the biosphere [22]. Specifically, we consider planetary habitability to be some function that is optimised when all of these elements are of some high value and no single element is dominant over or dominated by the others. Maximising only one of these (e.g. increased productivity through algal blooms) is not sufficient to enhance planetary habitability. Similarly, high resilience alone does not imply greater habitability – high resilience implies a systemic resistance to transformation or change of state (e.g. [23]). In the language of social and ecological resilience, a state has undesirable high resilience [23] where it has low diversity, low productivity, and low biomass locked-in for long intervals.

Here we attempt to enhance the utility of datasets from Earth’s deep history to provide comparison with contemporary biosphere change. For 31 episodes of macroevolutionary change in the geological record including that of the Anthropocene (tables 1, S1–S5) we consider their proximate cause (i.e., each disrupting agent) and assess the duration (figure S1; tables S3–S4) and impact (figures 2 and S2; table S5) of each disruptor on the biosphere. By evaluating the magnitude of anthropogenic biosphere disruption (a) so far, (b) likely to be caused by 2100, and (c) potentially caused by 3100 (table S5), we make quantitative and narrative comparisons between anthropogenic biosphere change and episodes of biosphere disruption in Earth history. Our analysis provides a firm set of baselines and methodology by which to evaluate the pace and magnitude of contemporary biotic change. It can thus inform policy on biodiversity loss, especially Goal A of the Global Biodiversity Framework (GBF), which is to halt human-induced extinctions, protect and restore ecosystem integrity, and significantly reduce the extinction risk of all species [24]. The GBF vision recognises that humans benefit from a healthy and resilient biosphere, and GBF Goal B is for people to prosper with nature. As we discuss with the disruptors concept below, Earth history shows how disrupting agents have both enhanced and degraded the biosphere. Humans are a disrupting agent of the biosphere, and we can draw lessons from Earth history to seek pathways to attain multigenerational prosperity with nature.

**Figure 2.**
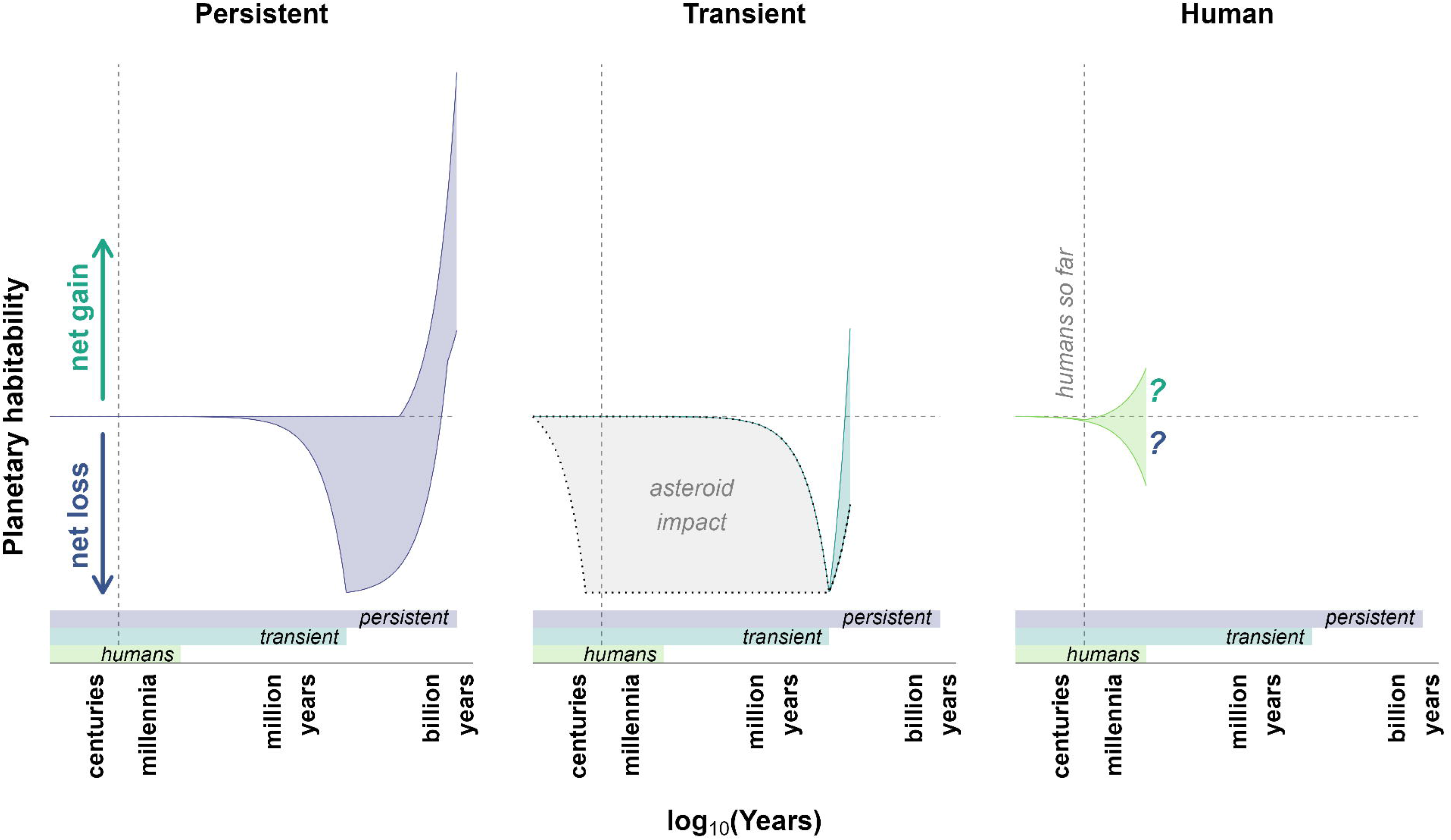
Schematic representation of the possible biosphere impacts from *persistent* and *transient disruptors* compared with human disruption. *Persistent disruptors* may initially cause biosphere degradation before overall biosphere growth as they persist in the Earth System. *Transient disruptors* drive biosphere degradation whilst present in the Earth System; biosphere recovery or transformation occurs after the disruptor and its effects dissipate. The y-axis scale is consistent across panels and represents proportional loss/gain; arrows are of equal length; horizontal dashed line represents no change; vertical dashed line shows the humanity-so-far interval; bottom bars represent disruptor durations; grey box shows the effect of an asteroid impact.

## 2. Biosphere disruptors

We resolve the driving mechanisms behind past episodes of biosphere change into two categories – *transient disruptors* (table S1) and *persistent disruptors* (table S2). *Transient disruptors* are geologically short-lived, ephemeral agents (figure S1). Our data show that *transient disruptors* tend to leave the biosphere with lower functional and taxonomic biodiversity, biomass, and productivity, locking in a metastable state of undesirable resilience [23] while the disruptor and its effects dissipate from the Earth System (figure S2). Biosphere recovery, reorganisation, and/or transformation to a new state of more desirable resilience typically takes several million years after the disruptor and its effects have dissipated (e.g. [25]). Indeed, the recovery profiles of different organism groups to transient disruptors are often complex, with some recovering rapidly, others more slowly, and with no simple relationship with the degree of biodiversity loss [26].

In contrast, *persistent disruptors* are geologically long-lasting and remain in the Earth System beyond the interval of their initial impact. An emergent feature of our data is that *persistent disruptors* have driven substantial increases in planetary habitability (functional and taxonomic biodiversity, biomass, productivity, and resilience) over Earth history (figure S2) by reconfiguring biosphere structure and function. However, *persistent disruptors* do not necessarily increase planetary habitability (figure 2). The Gaian bottleneck hypothesis asserts that life fails to persist on most planets where conditions are suitable for life to originate because planetary conditions move beyond the habitable range faster than life can evolve Gaian regulation to maintain habitability (e.g., through runaway greenhouse or icehouse climate states) [27]. In this context, the runaway climate state would be a *persistent disruptor*, locking in an undesirable biosphere state [23] of low (or no) biodiversity, productivity, and biomass [27]. Although *persistent disruptors* in Earth history have tended to increase planetary habitability, it is possible that humans could become a *persistent* but degradational disruptor of Earth’s biosphere (figures 2 and S2).

Uniquely as an agent of change in Earth’s history, humans have the *capacity to reflect on, govern, and potentially consciously modify our impacts* on the Earth System. Are we necessarily a *transient disruptor* of the biosphere, causing significant but ultimately short-lived losses of functional and taxonomic biodiversity and biomass [28]? Or do we have the potential to become a *persistent disruptor*, facilitating a transformation of the biosphere into a new configuration [29] that drives sustained enhancement of planetary habitability over longer time scales?

## 3. Methods

We identified 31 episodes of macroevolutionary change throughout Earth history that are (a) recognisable in the geological record including that of the Anthropocene, and (b) global rather than regional in extent and impact. Macroevolutionary episodes include radiations (increases in functional and taxonomic diversity), extinctions (decreases in functional and taxonomic diversity), and turnovers (changes in taxonomic composition with or without a net biodiversity change). These episodes of macroevolutionary change are detailed in tables S1–S5 and key episodes are outlined in table 1.

**Table 1.**
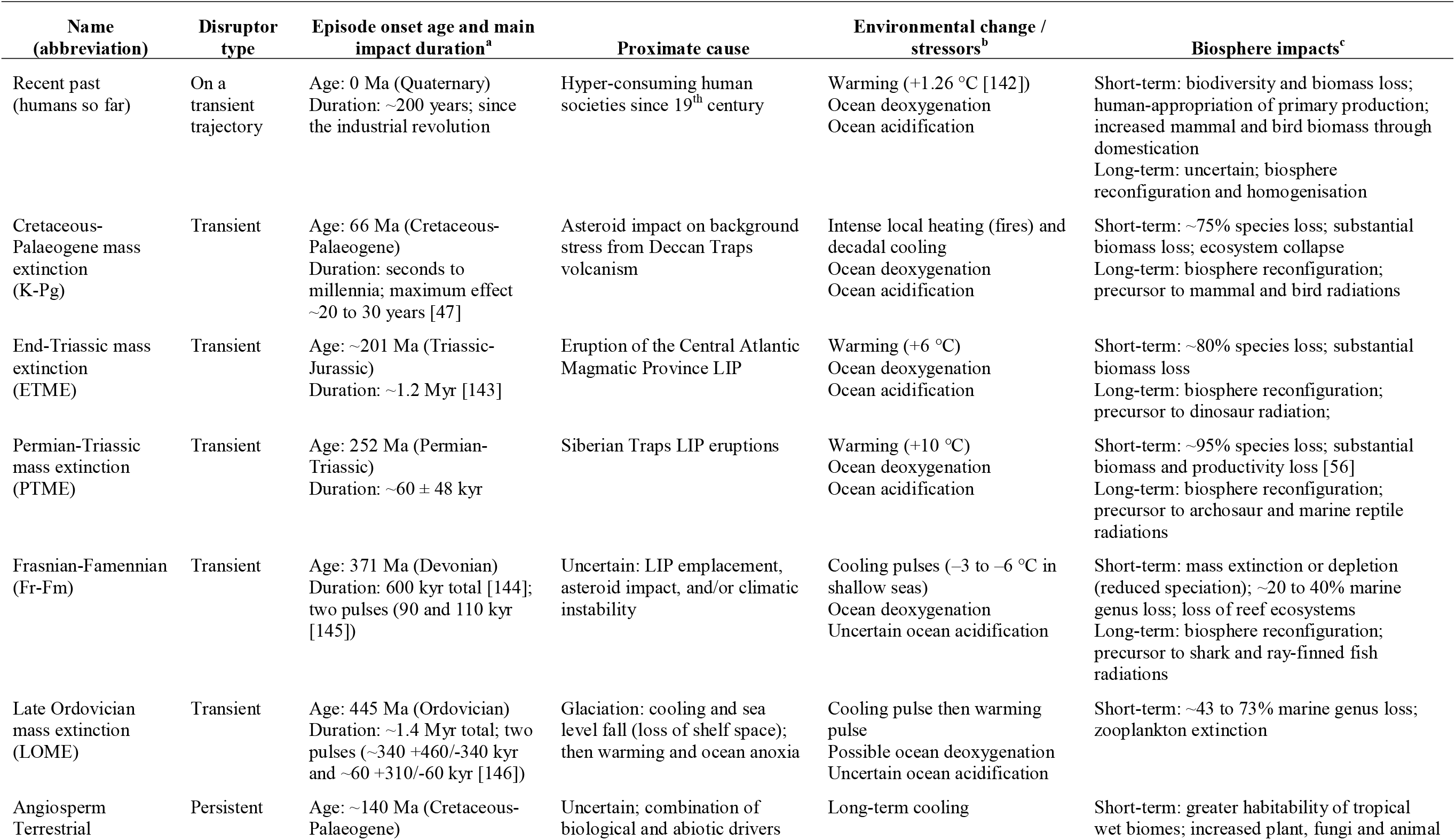
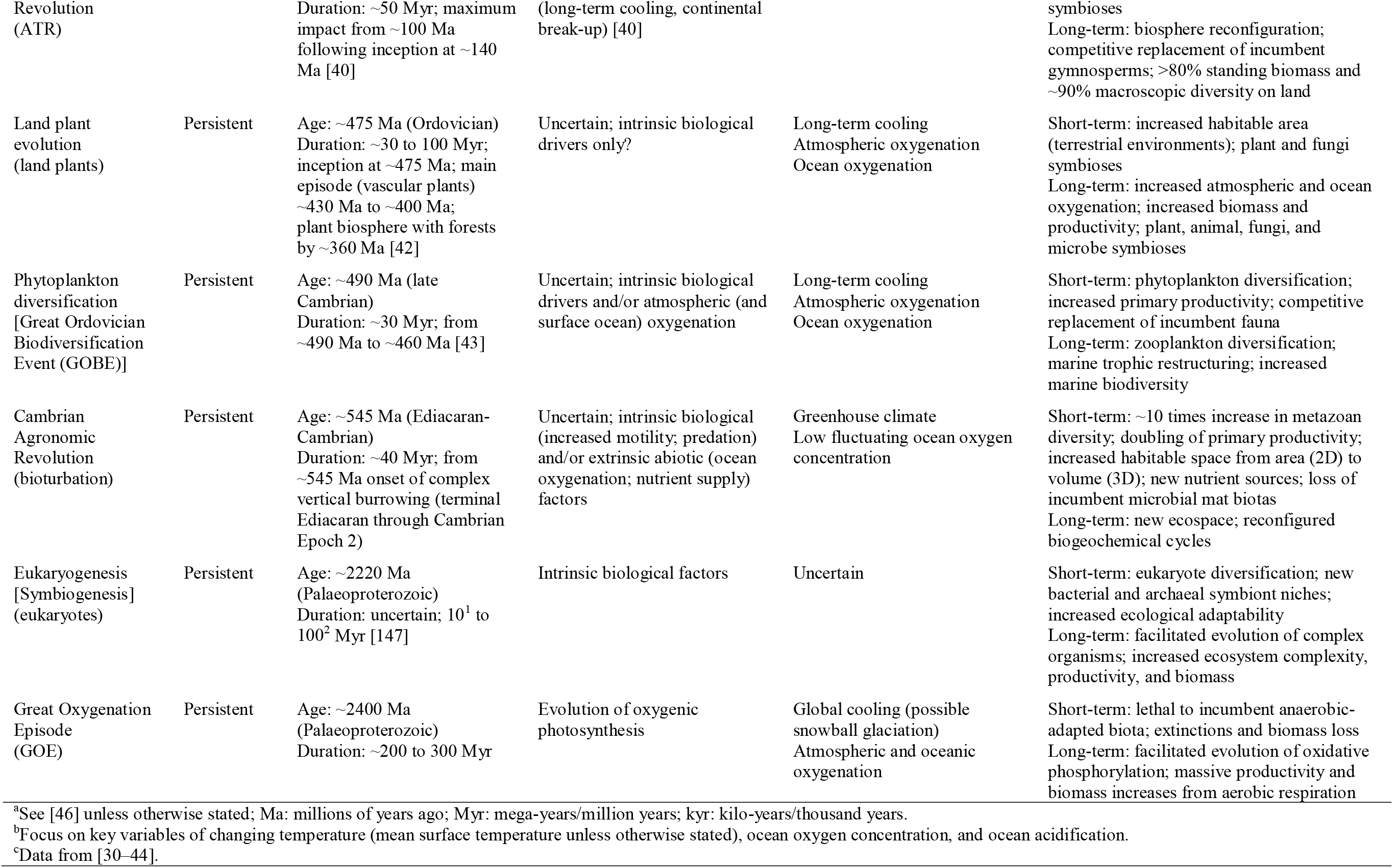
A selection of macroevolutionary episodes evaluated in this study: humans’ recent activity, the ‘Big 5’ mass extinctions, and the six quantified persistent disruptors. Macroevolutionary episodes are intervals of pronounced biodiversity gain (radiation), loss (extinction), or change (turnover), or of major biomass or productivity changes, at a global scale and that are evident in the geological record. Full dataset in tables S1 to S5.

Relevant literature on past biotic crises is well compiled elsewhere, and we ground our analysis of these episodes in review papers on biosphere change over eon or era timescales [30–38], along with primary literature searches for specific events that were not well represented in these reviews (tables S1, S3, S5). Episodes of macroevolutionary change that enhanced planetary habitability were selected because of their signals of increased adaptability, productivity, biomass, and biodiversity in the fossil record. In contrast to intervals of biotic crisis, there is a paucity of reviews quantifying intervals of biosphere growth across the Phanerozoic. We therefore relied on papers reviewing specific episodes of biosphere expansion (e.g. [39–44]), and reviews of the long-term evolution of biomass [37] and productivity [38] (tables 1, S2–S3, S5). We also note macroevolutionary changes that do not have a clear expression in the geological and/or fossil records, including gene transfer, RuBisCo, and Eukaryogeneisis (symbiogenesis); we do not attempt to quantify the biosphere impact of these episodes but include a narrative analysis of them (table S2).

For the 31 episodes of macroevolutionary change in the geological record, we evaluated their duration (figure S1; tables S3–S4), and magnitude (figure S2; table S5) and rate (figure 3; table S5) of biosphere change. We evaluated anthropogenic biosphere disruption (a) likely to be caused by 2100 and (b) potentially caused by 3100; i.e., on near-future human timescales (tables S1–S3, S5). We used the International Chronostratigraphic Chart v2024-12 [45] and The Geological Time Scale 2020 [46] for the ages of disruption episodes. We used primary literature searches to identify more precise age constraints for individual episodes where available (tables S1–3).

**Figure 3.**
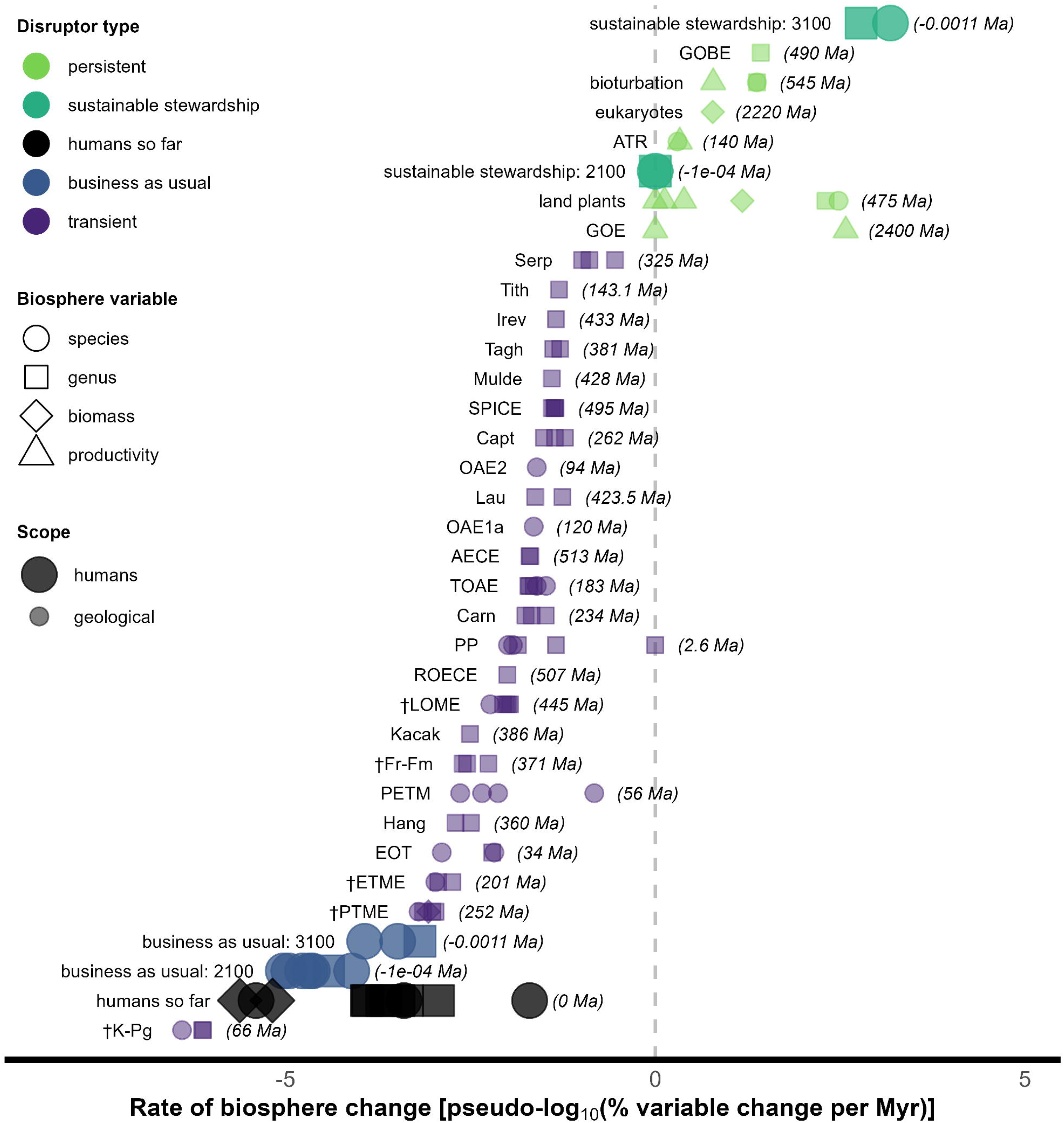
Humanity is driving the second highest rate of negative biosphere change in Earth history, after the K-Pg asteroid impact, but we could become the greatest driver of positive biosphere change over the next millennium. Rates of biosphere disruption (human: large symbols; Earth history: small symbols) as percentage biosphere variable change per duration in Myr on a pseudo-log_10_ scale (changes smaller than ±1% are treated as zero). Dagger-marks (†) indicate the ‘Big 5’ mass extinctions. The age of each episode is in parentheses on the righthand side. Data in tables S3 and S5.

To compare ancient, recent, and near-future biosphere change we evaluated percentage changes across episodes for species richness, genus richness, biomass, and productivity, where data were available. Data were drawn from across the palaeobiological and ecological literature (table S5), with the review papers referenced above used as the main sources of ancient macroevolutionary change data. Additional primary literature searches were conducted for episodes where data were sparse or that were not included in previous syntheses.

Biodiversity changes in palaeontological literature are typically reported as percentage species (or genus) change. To facilitate comparison with anthropogenic biosphere change literature, we converted species (or genus) extinction or origination percentage changes to extinctions or originations (E or O) per million species (genus) years (MSY [MGY]) following equation (1).

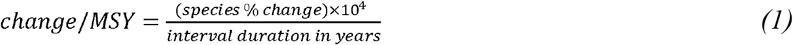

Percentage biosphere changes are presented on pseudo-log_10_ scales following equation (2). This allows both biosphere enhancement (positive) and degradation (negative) values to be represented on a consistent log scale, with negligible changes represented as zeroes.

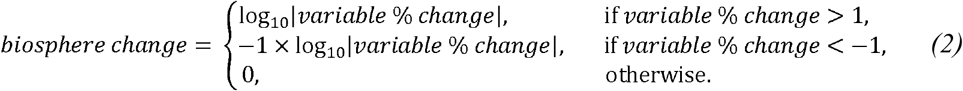

## 4. Ancient comparators for human activity

No past disruptors are directly comparable to contemporary biosphere change. This is partly due to the rapidity of anthropogenic changes – only the Cretaceous-Paleogene (K-Pg) asteroid impact is likely to have had its maximum impact over a shorter timescale than humans (figure S1; e.g. [47]). It is also partly because of the unique characteristics of the Earth System before and during each past episode of biosphere disruption, such as elevated early Palaeozoic extinction intensity being linked to lower atmospheric and oceanic oxygen concentration at the time [48].

Nevertheless, the *degree* of contemporary biosphere change can be compared to changes in deep time. We find that humans are degrading the wild biosphere with comparable magnitude to past transient biosphere disruptors (figure S2). We also find that humans have the potential to deliver the greatest rates of positive or negative change on the biosphere of all past disruptors, except for the K-Pg asteroid impact (figure 3).

### (a) Transient disruptors

*Transient disruptors* are geologically short-lived (figure S1; tables S1, S3–S4), with the disrupting force dissipating over time, even if the disruption caused to the biosphere continues long afterwards. In our dataset, *transient disruptors* have a mean duration of 8.0×10^5^ years (standard deviation 8.8×10^5^ years; median 5.0×10^5^ years; n = 25; figure S1; tables S1, S3–S4).

*Transient disruptors* are generally abiotic agents, including asteroid impacts and solid Earth processes (table S1). Although many solid Earth processes operate over long timescales (10^7^ to 10^8^ years), they also express as shorter duration pulses (10^4^ to 10^6^ years) that can be experienced as critical moments or tipping points in environmental systems that cascade through the biosphere (e.g. [49–51]). Volcanic outgassing occurs over tens of millions of years, but large igneous provinces (LIPs) can develop rapidly from mantle plumes, delivering climate-altering gases and aerosols into the atmosphere on shorter timescales (often <2×10^6^ years) with much of the outgassing taking place in even shorter pulses (10^4^ to 10^6^ years) [51]. Tectonic plates also move slowly, on timescales of 10^7^ to 10^9^ years, but the biosphere experiences crucial moments over shorter timescales (10^6^ to 10^7^ years) when changes in plate positions cause effective separation or connection of land masses and oceans. Changing continental configurations can also trigger substantial regional changes in biodiversity (e.g., the Great American Biotic Interchange [52]), but because these are not global-scale biosphere disruptions we do not include them here. More rarely, climatically driven changes influenced by the biosphere may have driven extinctions (e.g., with the Devonian expansion of land plants [53], though the relative impact is debated [54]).

*Transient disruptors* in Earth history have typically degraded the biosphere (figures 3 and S2), with extinctions resulting from mechanisms including hypoxia, excess CO_2_ (hypercapnia, ocean acidification, and acid rain), and rapid temperature change with coupled climate impacts such as aridity [33,35]. *Transient disruptors* generally result in lower biodiversity, simpler trophic structures [25,55], and reduced biomass [37,56] (figures 3 and S2). The biosphere then enters survival and recovery phases, often lasting millions of years, during which further extinctions might occur (e.g. [25]) before a new biodiverse biosphere re-evolves in the absence of the *transient disruptor*. Although the re-emerged biosphere may ultimately become more diverse than the one that went extinct, this happens once the effects of the *transient disruptor* have dissipated (e.g., after the K-Pg asteroid impact).

During the Permian-Triassic Mass Extinction (PTME), deforestation was widespread, particularly at low latitudes [56], probably caused by a combination of global warming, aridity, and acid rain (e.g. [33]). Net primary productivity on land was estimated to be ∼70% lower in the earliest Triassic (∼13.0 to 19.7 Pg C/yr) than in the latest Permian (∼54.4 to 62.5 Pg C/yr) [56]. This loss of primary productivity and the loss of genetic and functional diversity [25,57] threatened the biosphere integrity boundary in the PTME, just as contemporary biosphere integrity is threatened by loss of genetic diversity and HANPP (figure 1) [3]. By comparison, about one-third of global forest area has been lost since the Middle Holocene, with half of that loss occurring since 1900 and the most recent losses focused on the tropics [58]. Current HANPP is estimated at ∼30% (16.8 Pg C/yr) of the mean Holocene value [3]. Anthropogenic deforestation is approximately half that of the PTME, and the loss of undocumented biodiversity in tropical forests may already have crossed the mass extinction threshold [59].

Increased temperatures and expanded low-oxygen conditions in marine environments are a predominant feature of past biotic crises (table S1) [33,35]. Model simulations suggest that the combined effects of ocean deoxygenation (∼80%) and increased sea surface temperature (+10°C) can replicate ∼50% of PTME regional extinctions [60]. Marine hypoxia has not been accorded the same relevance as related patterns of climate change in driving current biodiversity loss [61,62]. Approximately 2% of the open ocean’s oxygen has been lost since 1960 [63], with a 3 to 4% loss predicted by 2100, and potential widespread marine oxygen deficiency within a millennium ([62] and references therein). Coastal hypoxia correlates with regions where the human footprint is strongest, linked to excessive nutrient flux and eutrophication [64].

Increasing ocean temperature reduces oxygen solubility (e.g. [63]), and there is empirical and theoretical evidence that a large increase in ocean temperature can disrupt phytoplankton photosynthesis [65,66], further disrupting the oxygen cycle. In some near-future scenarios where global warming reaches +4.9 ± 1.4°C by 2100, and +10 to 18°C by 2300 (relative to the 1850–1900 average), the combined effects of warming and oxygen depletion could result in extinction rates of comparable magnitude to previous mass extinctions, but limiting global warming to +2°C could reduce this risk by 70% [67].

Declining ocean oxygen concentration is a possible tipping element of global climate [68] with the potential to lead to a cascade effect throughout the Earth System [69]. However, ocean oxygen concentration is not explicitly included in the planetary boundaries framework; the phosphorus control variable is instead used as a proxy for the risk of ocean anoxia [2,3]. Ocean oxygen concentration is simpler to estimate in deep time than is phosphorus concentration, and dissolved oxygen has been measurably declining since at least 1960 [63,64]. Given the current trajectory of ocean oxygenation [62–64] and the association between biotic crises and ocean hypoxia in deep time (table S1) [33,35], we reinforce the suggestion of [61] that ocean oxygen concentration should be included explicitly in the planetary boundaries framework.

### (b) Persistent disruptors

In contrast to *transient disruptors*, *persistent disruptors* act on multi-million-year timescales (figure S1; tables S2–S4) and remain in the Earth System long after their initial impact. Here, we consider their duration as the interval over which their initial impact was greatest (range of best-estimate durations: 3.0×10^7^ to 5.0×10^8^ years; arithmetic mean ∼1.3×10^8^ years; standard deviation ∼1.7×10^8^ years, median ∼6.8×10^7^ years; n = 6; tables S3–S4).

In Earth history, *persistent disruptors* have typically promoted greater planetary habitability by supporting greater taxonomic and functional biodiversity, productivity, and biomass (figure S2) through new symbioses and interactions, even if they initially cause harm to the incumbent biosphere (figure 2). Persistence is not in itself desirable, and biologically impoverished (undesirable) system states can be resilient [23], but the *persistent disruptors* we have identified in the geological record have all driven substantial increases in planetary habitability (figures 3 and S2). *Persistent disruptors* primarily originate from within the biosphere itself (table S2), rather than from solid Earth processes. In this context, anthropogenic impacts on the biosphere also originate from within the biosphere.

The evolution of oxygenic photosynthesis is one of the earliest *persistent disruptors* evident in the geological record, known as the Great Oxygenation Episode (GOE) which began ∼2400 Ma and lasted ∼200 to 300 million years ([49] and references therein]. Oxygenic photosynthesis in the microbial biosphere caused the oxygenation of Earth’s surface environments and facilitated the subsequent diversity of more complex organisms, including eukaryotes, with higher energy demands satisfied by the greater energy released by oxidative phosphorylation [70]. However, the GOE caused significant harm to the incumbent biosphere that was adapted to oxygen-poor environments and predicated on anaerobic respiration pathways [71]. Primary productivity increased during the GOE, but fell back to a lower level for most of the Proterozoic [38]. Subsequent increases in atmospheric and oceanic oxygen concentrations from photosynthetic eukaryotes in the Mesoproterozoic and land plants in the Phanerozoic both further elevated primary productivity [38].

The evolution of land plants and the development of a terrestrial biosphere from ∼475 Ma [72] persistently disrupted the biosphere, fostering complex interactions between plants and fungi evident in some of the earliest terrestrial ecosystems [73]. Whilst newly evolved plants provided fungi with carbohydrates, fungi extended the soil volume from which plants could extract nutrients [74]. Animals then became key parts of these ecosystems from the Silurian and Devonian (e.g. [75]).

The late Mesozoic and early Cenozoic angiosperm (flowering plant) revolution may be seen as a further *persistent disruptor* embedded within the evolution of the terrestrial biosphere. This revolution drove terrestrial biodiversity and biomass to exceed its marine counterpart (table S2) [40] so that the terrestrial realm today hosts ∼85 to 95% of macroscopic species diversity [76]. The plant-dominated terrestrial biomass of 470 Pg C is almost two orders of magnitude greater than the marine biomass at 6 Pg C, despite similar rates of terrestrial and marine primary production (mass of carbon produced per unit time), with a greater turnover in the marine realm [77]. Angiosperms are the dominant component of terrestrial biomass and have higher productivity than gymnosperms [40]. The rise of angiosperms probably drove the decline in conifer diversity and their concomitant increased extinction risk [78]. Angiosperms evolved many mutualistic relationships with fungi and animals, with cascade effects massively enhancing the habitability of terrestrial tropical wet biomes [40]. The angiosperm radiation enabled the evolution of entire ecosystems that were previously impossible, and enhanced planetary habitability even whilst the incumbent conifer ecosystems declined and shifted into higher latitude, higher elevation, and lower productivity parts of the terrestrial realm.

## 5. Do some patterns of human interaction with the biosphere resemble persistent disruption?

Both *transient* and *persistent disruptors* permanently reshape the biosphere. *Transient disruptors* clear out biodiversity and break down existing trophic structures through (mass) extinctions, indirectly facilitating the permanent reshaping of the biosphere through the eventual evolution of new species and ecosystems. *Persistent disruptors* can reconfigure the state of the biosphere by promoting productivity, enhancing symbioses and mutualisms, and facilitating the evolution of new species and ecosystems.

Will humanity’s permanent reshaping of the biosphere (e.g. [79]) result from us acting as a *transient disruptor*, followed by a long interval of low functional and taxonomic diversity? Or can we identify pathways for humanity to transform the biosphere as a *persistent disruptor* that directly enhances global biodiversity? The latter scenario would seek to emulate the ‘Green Road’ shared socio-economic pathway SSP1 of greater equality, human well-being, lower overall consumption and growth, and greater sustainability [80]. However, the path of humans as a *transient disruptor* would run closer to the ‘Rocky Road’ SSP3 or ‘Highway’ SSP5 scenarios of high global environmental degradation, even if technology and conservation efforts prove effective locally [80]. Here we attempt to plot humans as both *transient* (business as usual; SSP3 or SSP5) and *persistent disruptors* (a more desirable social-ecological reorganisation; SSP1) in figure 3 (also figures 2 and S2; tables S3 and S5).

### (a) Fostering more diverse human influenced ecologies

Similar to *persistent disruptors*, some human practices increase the range of habitats available to life and foster more complex and diverse ecosystems, in part by leveraging local knowledge and community-based approaches to landscapes that develop and support new symbioses and consequently increase biodiversity. Many social-ecological systems support both human and non-human species around the world [15]. Indigenous communities around the North Pacific rim have maintained sustainable salmon fisheries for thousands of years by emphasising mutualism and multi-generational sustainability rather than maximising short-term exploitation [18].

In the Western Desert of Australia, human-landscape co-evolution over ∼50 kyr has enhanced biodiversity through processes of patch-mosaic burning that increased the number of ecological niches and reduced the risk of large destructive fires [17,81]. Winter burn-back of grassland increases the ecological space for other plants (e.g., bush tomatoes [81]), and the increased plant diversity supports more diverse and abundant lizard and mammal populations [17,82]. Patch-mosaic burning processes were developed by the Indigenous peoples of Australia to improve foraging and hunting outcomes [81]. Increased wild biodiversity and population sizes are an emergent property of this practice, which is rooted in societal values of social and environmental sustainability, and demonstrates that humans can act as biodiversity-enhancing ecosystem engineers on multi-millennial timescales [81]. Similarly, around the Mediterranean Basin, cork-oak savannahs support biodiverse mammal-, bird-, shrub-, and grassland ecologies [15] that have resulted from forest clearance, including by fire and livestock grazing, over millennia.

In monocultural industrial farming, ecological structure and diversity is much reduced. However, in many agricultural landscapes, both the range of crop types and their arrangement may increase wild biodiversity. Small-scale heterogeneity (within-farm or within-field for regenerative agriculture and agroforestry) can support greater invertebrate biodiversity, with larger scales needed to support enhanced vertebrate richness [83]. There is no one-size-fits-all approach, and a range of heterogeneities is needed to optimally support biodiversity [83], possibly with peak diversity achieved with a medium degree of heterogeneity [84].

Matrix farming intercalates areas of least impacted natural biosphere with mixed farmland uses that promotes biodiversity, species movement, and species interaction [16]. Such ‘working lands’ support local human communities and allow elements of the wild biosphere to move through the landscape by emphasising multifunctionality rather than maximising production of one particular animal or crop [16]. Conserving biodiversity in working landscapes will be essential to supporting wildlife in protected areas, where conservation areas alone may be too small to preserve biodiversity, especially as climate changes rapidly [16].

### (b) Sparing and sharing, land and sea

*Persistent disruptors* have improved biomass production, capture, and recycling, thereby providing a sustained increase in the energy available to the biosphere (table S5). Humans have increased total mammal biomass approximately threefold since 1850, but this is due to higher numbers and greater weights of domesticated livestock, and wild mammal biomass has approximately halved over the same time [13]. The human appropriation of mammal biomass and the primary productivity needed to sustain it has contributed to humanity driving the biosphere beyond its safe operating limits (figure 1) [1,3].

Conservationists have identified two major pathways to restoring biodiversity: ‘land sparing’, establishing separate areas for intensive agriculture and conservation [85]; and ‘land sharing’, developing multifunctional land use [86] where agricultural, nature restoration, and climate objectives are pursued within the same area. Humans may be approaching peak agriculture when we can begin to yield land back to wildlife [87], but this pattern is regionally complex, with some areas still experiencing agricultural expansion [88]. A shift from extensification to intensification of land use [88] may involve a combination of more diverse farming methods, including using a wider variety of plants [89], particularly those more resilient to climate change [89]), and combining ecological intensification with agro-ecological farming [90]. New technologies, including vertical farming systems, can increase agricultural productivity and reduce demand for land [91], as can increasing the photosynthetic efficiency of plants, especially those using C4 photosynthetic pathways [92]. Other agricultural innovations can improve and better maintain soil health and productivity, such as extending the symbioses between bacteria and plants to enhance nitrogen fixation [93]. In this new phase, land currently used for agriculture could be spared where suitable and sustained management could help biodiversity recovery alongside carbon capture [88].

However, the challenges are manifold. Agrarian labour constraints present a challenge to adapting practices towards systems that support higher biodiversity, and government subsidies may be required to make farming profitable while intensifying operations [94]. Moreover, the world needs at least a 50% increase in food production by 2050 to feed the projected 9.8 billion people, with demand for animal-based food rising by 70% [95]. Water and changing climate are serious limitations for agricultural innovations globally. If global reliance on animal-based foods declined from 20% to 5%, everyone could enjoy adequate nutrition while avoiding water deficits [96]. Climate change is reducing agricultural productivity in both calories and nutrients: every degree of heating reduces calorie production of staple crops per person per day by ∼120 kilocalories (∼4% recommended daily amount) [97]. Even accounting for adaptation measures, the impact of rising temperatures on production of staple crops remains negative and consequential [97]. If net zero carbon emissions can be reached quickly, global crop yields will decline by 11%, but if emissions rise unchecked, the decline is 24% [98]. More than one-third of all nations are self-sufficient in two or fewer of the seven food groups necessary for a healthy diet, making them vulnerable to supply chain disruptions [99]. Currently, more than two billion people suffer from malnutrition due to micronutrient insufficiency [100]. Managing the land with the twin goals of supplying adequate nutrition for people and bolstering biodiversity can be accomplished through a balance between physical and social constraints once the challenge is framed as transitioning to being a *persistent disruptor* of the land.

Biodiversity may not automatically recover following land abandonment (e.g. [101], and many landscapes will need input from humans for their restoration. A targeted approach to ecosystem restoration could limit further biodiversity loss, whilst also sequestering large volumes of carbon dioxide [102]. Many approaches towards ecosystem management and nature restoration, including rewilding [103] and policy measures to halt and reverse biodiversity decline (e.g. [24]) create co-benefits between climate mitigation from low-emission land use systems and nature recovery targets [104]. Humans have the potential to be positive forces of ecosystem engineering and biosphere enhancement where they maintain a persistent presence in landscapes and focus on multi-generational sustainability.

As with terrestrial ecosystems, the oceans reveal both the scale of human disruption and the scope for regenerative intervention. Although less than 10% of the global ocean is currently designated as a Marine Protected Area (MPA), and many marine ecoregions still have under 1% protection [105,106], recent governance developments suggest transformative expansion towards effective protection. Historically, targeted international interventions have demonstrated that marine ecosystems can rebound when coherent rules are applied. The 1911 North Pacific Fur Seal Treaty curtailed pelagic sealing, and later interwar bans on right whale hunting helped limit further depletion of vulnerable species [106]. These were precursors to the modern legal architecture: the 1982 United Nations Convention on the Law of the Sea (UNCLOS) [107], which establishes global duties to protect and preserve the marine environment, and the 1995 Fish Stocks Agreement (entered into force on 11^th^ December 2001) [108], which operationalizes precautionary and ecosystem-based approaches for straddling and highly migratory species.

Building on these foundations, the 2023 High Seas Biodiversity Treaty (BBNJ Agreement, entered into force on 17^th^ January 2026) [109] enables the creation of area-based management tools, including high seas MPAs, supported by environmental impact assessments and capacity-building measures. Overall, the BBNJ Agreement provides, for the first time, a mechanism to ‘spare’ portions of the high seas through legally established protected areas, a global stewardship complementing national exclusive economic zone (EEZ)-level stewardship. However, sparing marine volumes is practically more challenging than sparing land areas, as any boundaries put in place are more porous to deliberate and accidental incursion. It is important, therefore, to accompany specific partial protection measures for the ocean with global approaches that consider the interconnectedness of the whole marine realm.

At the interface of ecology and governance, ecosystem-based management has become central to reconciling seafood production with biodiversity conservation. Regional fisheries management organizations integrate ecological considerations into bycatch mitigation, habitat protection, and harvest strategies, though significant gaps in implementation remain. In parallel, emerging dynamic ocean management tools – informed by satellite data, species distribution models, and real-time environmental indicators – demonstrate that it is possible to reduce impacts on nontarget species while maintaining or even improving fisheries performance.

Taken together, these developments suggest that the oceans, like terrestrial systems, may be entering a phase where human presence – guided by coherent legal and policy frameworks, supported by science-based management – can increase the adaptive capacity and long-term transformability of marine social-ecological systems. Achieving the Kunming–Montreal Global Biodiversity Framework’s 30×30 target in the marine realm will require precisely this integration of spatial protection, effective governance, ecosystem-based management, and adaptive technological tools, to support marine biodiversity recovery while sustaining the food systems upon which billions (humans and non-humans) depend.

For humans to transition from a *transient* to *persistent disruptor* of the marine biosphere, a profound change in policy and legal approach will be necessary: from the prevalent ‘freedom of the seas’ [107,110] and the ‘common heritage of mankind’ ([107] Part XI) to the ‘responsibility for the seas’ approach [111,112]. This means transitioning from an extractive focus, characterising the ‘freedom of the seas’ and ‘common heritage of mankind’ principles, to a stewardship and protective approach (e.g. [113]). This shift illustrates how humans might move from acting as a *transient disruptor* of ocean systems to becoming *persistent* agents of enhanced planetary habitability, through stewardship, adaptive governance, and potentially transformation towards more desirable marine regimes.

This transition should not be expected to advance by moral or idealistic reasoning; rather it will be borne out of necessity. The Earth System, long implicitly considered a stable background for economic and political development through the Holocene, is increasingly understood (in the Anthropocene epoch) as simultaneously the precondition for human future, and vulnerable to disruption from overwhelming human impacts. Once this realisation is translated into political, economic/financial, and legal change of human (via states’) activity, humans may enshrine the prospects of evolving from a *transient* to a *persistent* disruptor. Ocean governance is a critical arena in which the broader transformation toward ‘common responsibility for the Earth System’ [114] must unfold. This can be driven internationally, as described above, as well as through national and corporate governance. Ecuador provided for the Earth System itself to have legal rights (the “Rights of Nature”) in the national constitution of 2008, which rights have been successfully used to defend the natural environment against corporate pollution, catastrophic water cycle changes, and increased species extinction risk [115]. At least 25 companies have made provision for the natural world to be represented in their corporate governance structures, and this representation may have a tangible impact on decision-making even in organisations whose mission are already nature-aligned [116].

### (c) Novel ecologies and associations of species

Land abandonment facilitates the emergence of novel ecologies that evolve after human activity has ceased. These communities might resemble recovery biotas like those following extinctions in deep time [25], but near-future recovery biotas would be uniquely different in that they will comprise a community with a more globally homogeneous complement of species through non-native translocations [117]. Novel terrestrial ecosystems are now typified by non-native species [118]. In many marine settings, non-native species currently play a smaller but growing role (especially on islands), and novel ecologies are characterised by range shifts and the unequal responses of species to environmental change [119]. In some coastal marine ecologies, the complement of species has been radically changed by species translocation, leading to the emergence of ecosystems with very different compositions [120]. Non-native species translocation is one of the major causes of biodiversity loss in ecosystems across the world (e.g. [121]). However, many non-native species merge into the native community without negative effects [122], thereby enriching rather than degrading the local ecosystem (e.g., species introductions have approximately doubled UK floral diversity [123]).

Although species extinction and reduced populations of wild species at the global level are now serious problems [28,124], ‘novel’ associations of species at the local and regional level (e.g., in urban areas or abandoned agricultural land) provide habitats to new or previously rare species through niche creation and mutualisms [125,126]. Processes of species translocation, agriculture and horticulture lead to more rapid hybridization and speciation, often within a few generations [20,127]. Hybridization is not well studied for marine ecologies, but examples include those between the Mediterranean mussel *Mytilus galloprovincialis* and the Pacific *Mytilus trossulus* [128]. On land, Thomas [20] noted ∼6 to 8 new hybrid species of cultivated crops resulting from human intervention and observed that most new agricultural and horticultural species are obligate mutualists with humans. Whilst many of the newly naturalised species are facultative mutualists, land plant speciation rates may now be the highest in their evolutionary history [20]. In co-opting humans to disperse their seeds, such patterns resemble the symbioses that formed between animals and flowering plants during the angiosperm revolution (table S2). There is also evidence for reproductive isolation leading to allopatric speciation in non-native species that are detached from the gene pools in their native range [21,129], likely leading to the emergence of neo-species. Non-native species can also drive rapid evolution in native populations as a response to competition [130], or through enhanced reproduction [131]. Though clearly noting the loss of irreplaceable populations and species, Thomas [132] suggested that processes of hybridization and allopatric speciation might plausibly lead to a net increase in global biodiversity. Nevertheless, species resulting from novel human ecologies should not be seen as a trade-off against the ones made extinct [126,133], and currently increased human activity is associated with decreased natural vegetation diversity globally [21,134].

Humans produce waste materials in vast quantities. Increasing volumes of this waste are hard-to-degrade plastics [135] that are harmful to the biosphere and pose significant environmental and human health challenges. However, the biosphere may offer some remediation. Both terrestrial [136,137] and marine [138] fungi and bacteria are able to break down many plastic-forming polymers. If new microbial ecologies can be fostered in areas of concentrated human-produced waste, there is potential both to support novel microbial communities and to reduce our environmental impacts with the support of the biosphere. The evolution of lignin decomposition which enabled the decay and recycling of woody tissues would represent a precedent in Earth history [139].

Novel assemblages, commonly with locally enhanced biodiversity, may become significant as climate change, the continued translocation of non-native species, and widespread toxic pollutants increasingly impact local assemblages. Such novel assemblages may serve, often inadvertently, as ‘environing technologies’, shaping landscapes and environments on micro, local, and planetary scales [140]. As the Anthropocene epoch unfolds, it may be that these assemblages emerge as the survivors within a progressively human-influenced global environment [141].

## 6. Conclusions

Humans have the capacity to recognise, govern, and potentially transform their impacts on the Earth System, even though the scale of this capacity is uneven and contested. Humans sometimes emulate *persistent disruptors* of the biosphere from Earth history which led to enhanced biosphere productivity, biomass, biodiversity, and greater mutualisms and symbioses between species (table S2). These *persistent disruptors* ultimately drove greater planetary habitability. Even accepting that some novel and managed ecologies are emerging that resemble the impact of past *persistent disruptors*, it seems prudent to preserve existing biodiversity and not threaten its integrity in ways that precipitate a crisis resembling the impact of past *transient disruptors* [28], which is our current trajectory. Earth history tells us that the path of a *transient disruptor* is through extinctions which lead to a depauperate biosphere for millions of years, with highly degraded ecosystems and much-reduced biomass and biodiversity. It also tells us that extinctions can be highly selective, with some ecological and biological groups impacted significantly more than others (e.g. [36]). If we continue our current trajectory, humans will have the second highest rate of negative effect on the biosphere in Earth history, after only the K-Pg asteroid impact (figure 3). However, we have the potential to drive the highest rate of positive impact on the biosphere in Earth history, and to do so on human timescales of decades to millennia (figure 3; table S5). For this to become possible, a profound change in the political and legal framework – towards a ‘common responsibility for the Earth System’ as a component of international law [114] – is a prerequisite.

We have aimed to show that the palaeontological record, often used as a comparison with contemporary biosphere decline, also shows how planetary habitability has increased over long timescales. Innovations from within the biosphere, such as the mutualisms forged by flowering plants, have driven sustained enhancement of biodiversity, productivity, and biomass over billions of years. Our analysis speaks directly to Goal A of the Kunming-Montreal Global Biodiversity Framework that aims to preserve and restore ecosystems and biodiversity [24], providing baselines from the history of life on Earth against which we can assess the pace and magnitude of contemporary biosphere change. Our analysis highlights and provides evidence from Earth history of the importance of Goal B, to prosper with nature [24], recognising that humans on the trajectory of a *transient disruptor* are degrading the biosphere, including essential ecosystem functions. Humans have the potential to learn from Earth’s deep history, to prosper with nature as a transformative and *persistent disruptor* of the biosphere, and to ultimately enhance planetary habitability.

## Acknowledgements

TWWH and MW acknowledge support from the Leverhulme Trust, grant RPG-2022-233. HB was partially supported by UK Research and Innovation through the Biotechnology and Biological Sciences Research Council with contributions from the Department for Environment, Food & Rural Affairs, Department for Energy Security and Net Zero, and Scottish Government (Land Use for Net Zero, Nature and People Hub, grant number BB/Y008723/1), with participation of the Department of Agriculture, Environment and Rural Affairs of Northern Ireland and Welsh Government. MY was supported by the Research Grants Council of the Hong Kong Special Administrative Region, China (RFS2223-7S02; HKU 17306023; CityU 17304025). SP was partially supported by the European Space Agency under contract No. 4000146344/24/I-LR and by the EU Horizon programme through the OceanNexus project. We are grateful to the editor and two anonymous reviewers whose thoughtful comments helped us improve the clarity of this manuscript.

## Data and code availability

The code used to produce figure 1 is available on GitHub at: https://github.com/twwh01/planetary_boundaries_plots. The supplementary data and code necessary to reproduce all other figures and analyses are available on GitHub at: https://github.com/twwh01/disruptors.

## Ethics and approvals

No ethical approval, licences, or permits were required for this study. No AI systems were used in this work.

