## Supplementary Information for "Humans could become the greatest driver of biosphere net gain in Earth history, but are currently the second fastest driver of biosphere loss"

**List of Supplementary Materials**

### Explanation of the Supplementary Data

**Table S1.** Examples of transient disruptors from Earth's deep past to contemporary associated with substantial habitat loss caused, for example, by global changes in climate and low oxygen concentrations in the marine realm. Unless otherwise stated, data follow [S1–S3].

**Table S2.** Examples of persistent disruptors to the biosphere, from Hadean to contemporary times. Humans are included as potential persistent disruptors. Sources of information include [S4] for gene transfer, [S5] for RuBisCo, [S6] for symbiogenesis (eukaryogenesis), [S7] for sexual reproduction and references in text.

**Table S3.** The ages and duration of impact of transient and persistent disruptors in Earth history. These data are shown in figures S1-S2 and are used to calculate the rates of change in figure 3.

**Table S4.** Summary statistics of the event durations listed in table S3.

**Table S5.** The degree of change of four biosphere variables (species richness, genus richness, biomass, and productivity) through past transient and persistent disruption events and potential human futures. These data are shown in figure S1 and used to calculate the rates of change in figure 3.

### Supplementary Figures

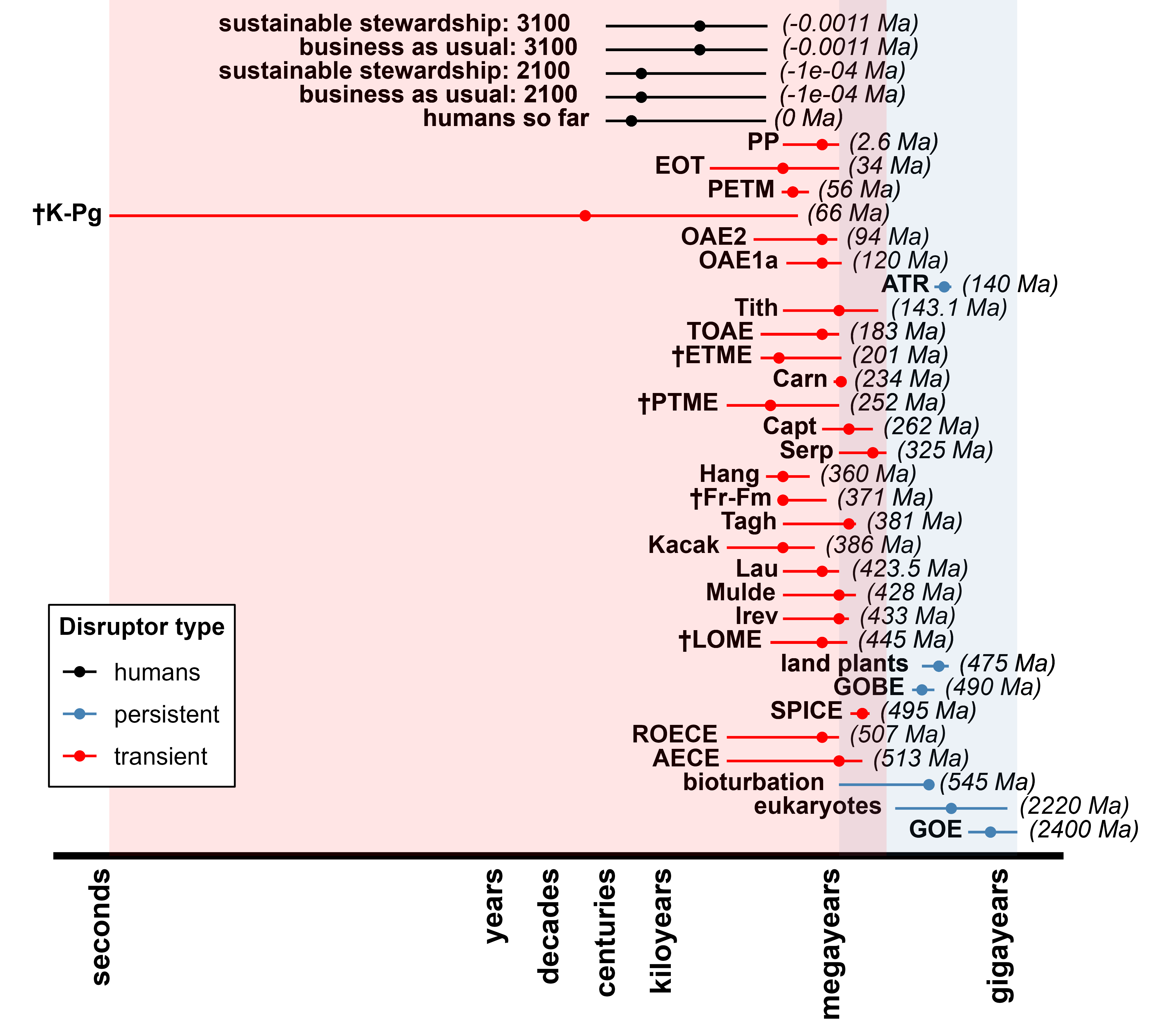

Figure S1. Timescales (log_10_) of *human disruption* (black) alongside *transient* (red) and *persistent* (blue) *disruptors* in Earth history. Each line represents the range of timescales over which the disruptor acted or will act on the biosphere with points indicating the duration of main impact. Dagger-marks (†) indicate the ‘Big 5’ mass extinctions. The age of each disruption in millions of years ago (Ma) is in parentheses. See table S3 for data and abbreviations.

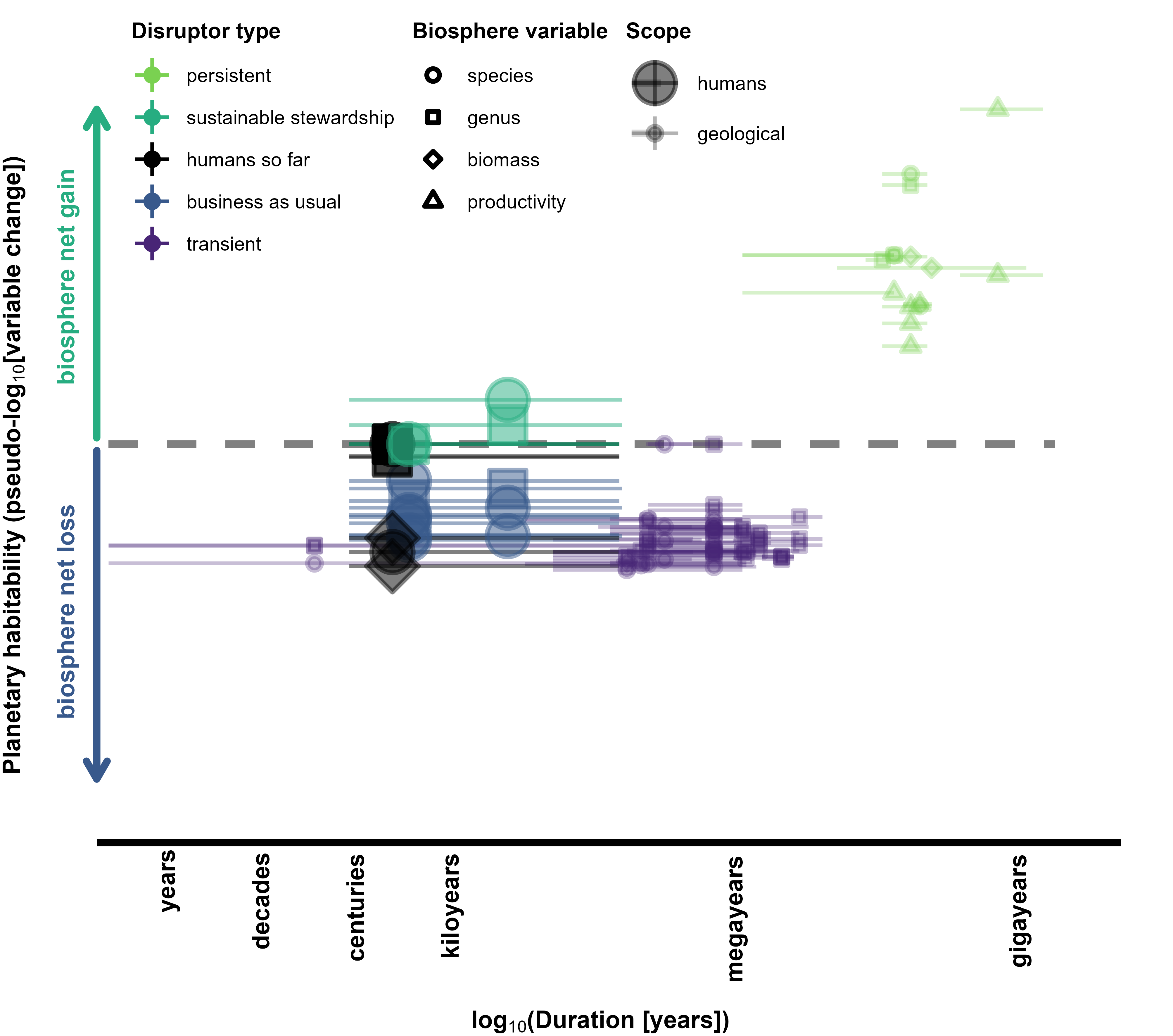

Figure S2. Timescales and impacts of human disruption (large symbols) alongside *transient* and *persistent disruptors* in Earth history (small symbols). Human activity plots with past *transient disruptors* for the recent past and business as usual near-future. *Persistent disruptors* from Earth history operated on long timescales and drove biosphere net gain. Human activity under a sustainable stewardship scenario has the potential to drive biosphere net gain. Axes are log_10_ scales. The y-axis shows percentage change (positive or negative) of biosphere variable (species richness, genus richness, biomass, and productivity) on a pseudo-log_10_ scale with percentages changes smaller than ±1% treated as zero. Points are plotted at their central duration estimate with error bars representing the duration range. Data in tables S3 and S5.
